# Disrupted Frontoparietal Dynamics in Neurofibromatosis Type 1: Reduced Sensitivity and Atypical Modulation During Working Memory

**DOI:** 10.1101/2025.04.10.648183

**Authors:** Marta C Litwińczuk, Shruti Garg, Caroline Lea-Carnall, Nelson J Trujillo-Barreto

**Affiliations:** School of Health Sciences, University of Manchester, Manchester, UK; Division of Psychology and Mental Health, Faculty of Biology, Medicine and Health, University of Manchester, Manchester, UK; Royal Manchester Children’s Hospital, Manchester University NHS Foundation Trust, Manchester, UK; Geoffrey Jefferson Brain Research Centre, Manchester Academic Health Science Centre, Manchester, United Kingdom; School of Psychology, Manchester Metropolitan University, Manchester, UK

**Keywords:** Neurofibromatosis, working memory, effective connectivity, functional MRI

## Abstract

Neurofibromatosis Type 1 (NF1) is a rare, single-gene neurodevelopmental disorder. Atypical brain activation patterns have been linked to working memory difficulties in individuals with NF1. The present work investigates if NF1 has increased inhibitory activity in the frontoparietal network during working memory tasks compared to neurotypical controls. Forty-three adolescents with NF1 and twenty-six age-matched neurotypical controls completed functional magnetic resonance imaging scans during a verbal working memory task. Dynamic causal models (DCMs) were estimated for bilateral frontoparietal network (dorsolateral and ventrolateral prefrontal cortices (dlPFC and vlPFC), superior and inferior parietal gyri (SPG and IPG)). The parametric empirical Bayes approach with Bayesian model reduction was used to test the hypothesis that NF1 diagnosis would be characterised by greater inhibitory self-connections (intrinsic connectivity). Leave-one-out cross-validation (LOO-CV) was performed to test the generalisability of group differences. NF1 participants demonstrated greater average intrinsic connectivity of left dlPFC, IPG, SPG and bilateral vlPFC. The DCM that best explained effects of working memory showed that NF1 group has increased intrinsic connectivity of left vlPFC, but weaker intrinsic connectivity of right vlPFC and left dlPFC. The parameters of these connections showed a modest but positive predictive correlation of r = 0.19 (p = 0.055) with diagnosis status, suggesting a trend toward predictive value. Overall, increased average intrinsic connectivity of left dlPFC, IPG, SPG and bilateral vlPFC in NF1, suggests reduced overall sensitivity of these regions to inputs. Working memory evoked different patterns of input processing in NF1, that cannot be characterised by increased inhibition alone. Instead, modulatory connectivity related to working memory showed less inhibitory self-connectivity of left dlPFC and left vlPFC, and more inhibitory intrinsic connectivity of right vlPFC in NF1. This discrepancy between average and modulatory connectivity suggests that overall NF1 participants are responsive to cognitive task-related inputs but may show atypical adaptation to the task demands of working memory.

## Introduction

Neurofibromatosis Type 1 (NF1) is a rare, single-gene autosomal dominant neurodevelopmental disorder with birth incidence of 1:2,700 (Evans et al., 2010). Affected individuals present with a variety of symptom expression and severity, including pigmentary lesions (café-au-lait spots), dermal neurofibromas, skeletal abnormalities, brain and peripheral nerve tumours, learning disabilities, and cognitive and social deficits (Gutmann et al., 2017). Individuals with NF1 also present with high rates of comorbidity with attention-deficit/hyperactivity disorder and Autism Spectrum Conditions (ASC) (Garg et al., 2013; Garg et al., 2012; Matson et al., 2013; van der Meer et al., 2012).

NF1 is caused by mutations to the NF1 gene, which encodes the neurofibromin protein. Mutations in NF1 gene lead to reduced or aberrant neurofibromin function and consequently to hyperactivation of the RasMAPK signalling pathway (Daston & Ratner, 2005; North et al., 1997). This dysregulation has downstream effects of increasing presynaptic gamma-aminobutyric acid (GABA) activity (Costa et al., 2002). Alterations in GABAergic activity is associated with disruption in neural development (van Lier et al., 2020). Additionally, GABA plays a crucial role in regulating neuronal activity, through selective suppression of firing of neurons (Bannai et al., 2015), and excessive GABA activity can disrupt the cortical excitation/inhibition balance (E/I) (Maffei et al., 2017). The E/I balance is essential for maintenance of function of central nervous system, and has further implication on regional processing of cognitive tasks, and communication of input-related signals between regions (Metkus et al., 2024; Sears & Hewett, 2021; Sohal & Rubenstein, 2019). Therefore, the selective effect of NF1 on the GABAergic activity has broad implications for understanding the role of E/I balance in the mechanisms of cognition and neurodevelopmental conditions.

NF1 presents with a complex profile of cognitive, behavioural and social disruptions. Research comparing NF1 individuals to their unaffected siblings documented academic underperformance and cognitive difficulties (Lehtonen et al., 2012), including the domains of general intelligence, visuospatial processing, executive function, attention, and social cognition. Research literature has reliably demonstrated working memory difficulties in NF1 (Lehtonen et al., 2012; Pobric et al., 2021; Sawyer et al., 2022; Shilyansky et al., 2010) which is the ability to temporarily maintain and manipulate information in one’s memory (Baddeley, 1992; Baddeley et al., 2014). Working memory underlies for our ability to perform complex tasks and it is critical for academic achievement, including language acquisition, reading comprehension and mathematics performance (Gathercole & Alloway, 2005). Consequently, the neural and cognitive mechanisms of working memory disruptions in NF1 may contribute to their broader academic and cognitive challenges.

Several neuroimaging studies have highlighted the effects of NF1 on neural function during working memory. In a seminal study, Shilyansky et al. (2010) conducted parallel neuroimaging research of working memory performance and learning in humans and mice with the NF1 mutation. Human research showed that compared to neurotypical controls, NF1 participants were less accurate and slower during performance of a working memory task. The authors also investigated the neural correlates of these disruptions in NF1 with functional magnetic resonance imaging (fMRI). Compared to neurotypical controls, NF1 group had weaker activation in parts of the distributed working memory network -including the dorsolateral prefrontal cortex (dlPFC), frontal eye fields, parietal cortex, and striatum. These findings paralleled with electrophysiological findings of elevated inhibitory activity in prefrontal cortical and striatal regions from Nf1+/-mouse models. Therefore, Shilyansky et al. (2010) proposed that these patterns of regional hypoactivation in human frontal and parietal regions may be directly linked to increased GABAergic inhibition. Ibrahim et al. (2017) extended these findings with investigation of communication within the frontoparietal network. By investigating functional connectivity during working memory performance with generalized psychophysiological interaction (gPPI) method, Ibrahim et al. (2017) demonstrated that in response to increased working memory load, the neurotypical control group had greater connectivity between the right parietal seed and critical working memory-related regions, including bilateral parietal cortex, frontal areas (right pars opercularis and left premotor cortex). This suggests that the individuals with NF1 may have difficulty in modulating their connectivity within the frontoparietal network in response to increasing working memory demands.

The next step to understanding the mechanisms of working memory performance in individuals affected by NF1 is to investigate how E/I balance may influence the patterns of connectivity within the frontoparietal network. Therefore, this work aims to investigate whether the NF1 group has increased inhibitory activity in the fronto-parietal network, as compared to control group. To achieve this, we applied dynamic causal modelling (DCM) to fMRI data collected during a verbal working memory task (N-back task) from adolescents diagnosed with NF1 and age-matched neurotypical controls. DCM is a framework that allows the estimation of directed interactions (both excitatory and inhibitory) between neuronal populations, known as effective connectivity (Zeidman, Jafarian, Corbin, et al., 2019). In addition and of particular interest to this work, DCM also estimates intrinsic connectivity, which reflects the self-inhibitory dynamics of a region and, by extension, its responsiveness (sensitivity) to external inputs (Snyder et al., 2021). Importantly, intrinsic connectivity serves as a proxy for the region’s E/I balance, which is regulated by GABAergic neurotransmission (Maffei et al., 2017). In the context of NF1, elevated GABA levels have been proposed to disrupt this balance (Buratti et al., 2012; Payne et al., 2010; Petroff, 2016). Based on this evidence, we hypothesize that the NF1 group will have increased inhibitory intrinsic connectivity in the frontoparietal network during N-back tasks compared to neurotypical controls.

## Methods

### NF1 Participants

Part of the data used here is from a previous study described in detail in Garg et al. (2022). Forty-three adolescents aged 11–17 years were recruited via the Northern UK NF-National Institute of Health, with (i) diagnostic criteria [National Institutes of Health Consensus Development Conference. Neurofibromatosis conference statement. Arch. Neurol. 45, 575–578 (1988).] and/or molecular diagnosis of NF1; (ii) no history of intracranial pathology other than asymptomatic optic pathway or other asymptomatic and untreated NF1-associated white matter lesion or glioma; (iii) no history of epilepsy or any major mental illness; and (iv) no MRI contraindications. Participants on pre-existing medications such as stimulants, melatonin or selective serotonin re-uptake inhibitors were not excluded from participation. The study was conducted in accordance with local ethics committee approval (Ethics reference: 18/NW/0762, ClinicalTrials.gov Identifier: NCT0499142. Registered 5th August 2021; retrospectively registered, https://clinicaltrials.gov/ct2/show/NCT04991428). All methods were carried out in accordance with relevant guidelines and regulations.

### Neurotypical controls

Twenty-six adolescents with no diagnosis of NF1 were recruited. Controls’ age and sex were matched to the sample of NF1 group. Table 1 includes demographics information for NF1 and control groups.

**Table 1.** Demographic data of study participants (43 NF1 participants, 26 neurotypical controls).

|  | Mean Age (STD) | Sex (M/F) |
| --- | --- | --- |
| NF1 | 14.8 (1.83) | 22/21 |
| Controls | 14.15 (2.71) | 12/14 |

### Experimental Procedure

Participants laid in the scanner and a T1-weighted image was acquired. Participants were asked to perform a working memory task for 6 min while fMRI scans were acquired. In addition, T2-weighted images were acquired at the first visit (after the T1 image) and reviewed by a paediatric neuro-radiologist to rule out NF1 associated tumours.

The working memory task session consisted of 6 blocks each of 0-back and 2-back verbal N-back task (Figure 1). During the 0-back condition, participants were instructed to press a hand-held button only when the letter ‘X’ was presented on the screen. For the 2-back condition, the participants were instructed to press the same button when the letter on the screen matched the letter 2 presentations before. Each block was 30 seconds long and consisted of 9 target stimuli.

**Figure 1.**
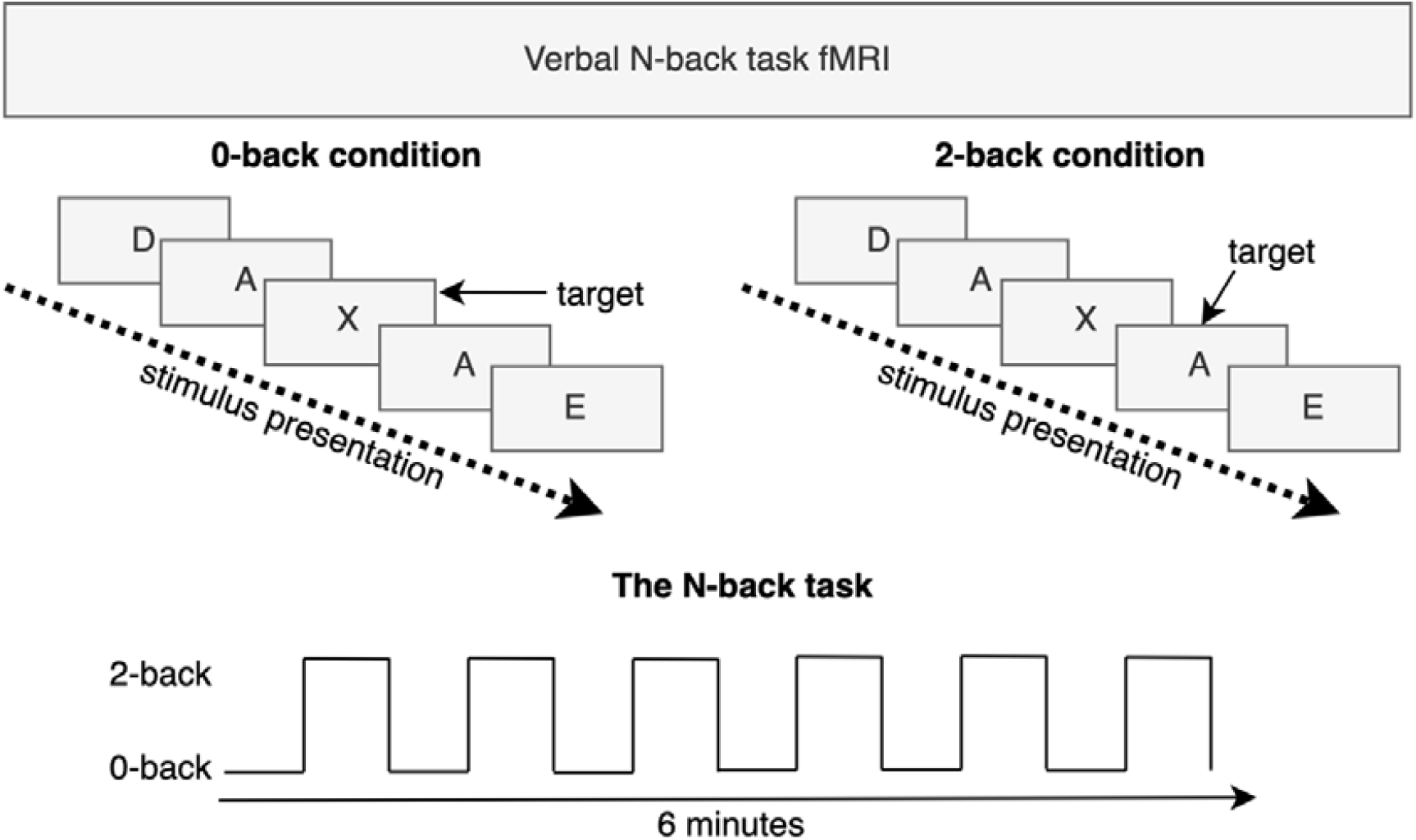
A visual illustration of the verbal working memory task that the participants performed during scanning.

### Image acquisition

Imaging was performed on a 3 Tesla Philips Achieva scanner using a 32-channel head coil with a SENSE factor 2.5. To maximise signal-to-noise (SNR), we utilised a dual-echo fMRI protocol developed by Halai et al. (2014). The fMRI sequence included 36 slices, 64×64 matrix, field of view (FOV) 224×126×224 mm, in-plane resolution 2.5×2.5 mm, slice thickness 3.5 mm, TR=2.5 s, TE = 12 ms and 35 ms. The total number of volumes collected for each fMRI run was 144.

### Image Processing

Image processing was done using SPM12 (Wellcome Department of Imaging Neuroscience, London; http://www.fil.ion.ucl.ac.uk/spm) and MATLAB R2023a. Dual echo images were extracted and averaged using in-house MATLAB code developed by Halai et al. (2014) (DEToolbox). First, functional images were slice-time corrected and realigned to first image. Then, the short and long echo times were combined for each timepoint. The orientation and location of origin point of every anatomical T1 image was checked and corrected where needed. Mean functional EPI image was co-registered to the structural (T1) image. Motion parameters estimated during co-registration of short images were used as input to the Artifact Detection Tools (ART; https://www.nitrc.org/projects/artifact_detect/) toolbox along with combined dual echo scans for identification of outlier and motion corrupted images across the complete scan. The outlier detection threshold was set to changes in global signal 3 standard deviations away from mean global brain activation. Motion threshold for identifying scans to be censored was set to 3 mm. Outlier images and images corrupted by motion were censored during the analysis by using the outlier volume regressors. Subjects with <80% of scans were removed from analysis. Following removal of participant data with high motion and artefacts, 26 controls and 43 NF1 participants remained. Unified segmentation was conducted to identify grey matter, white matter, and cerebrospinal fluid. Normalisation to MNI space was done with diffeomorphic anatomical registration using exponentiated lie-algebra (DARTEL) (Ashburner, 2007) registration method for fMRI. Normalised images were interpolated to isotropic 2.5 × 2.5 × 2.5 mm voxel resolution. A 6×6×6mm full width at half maximum (FWHM) Gaussian smoothing kernel was applied.

### fMRI analysis

The fMRI analysis and connectivity modelling pipeline has been visually summarised in Figure 2.

**Figure 2.**
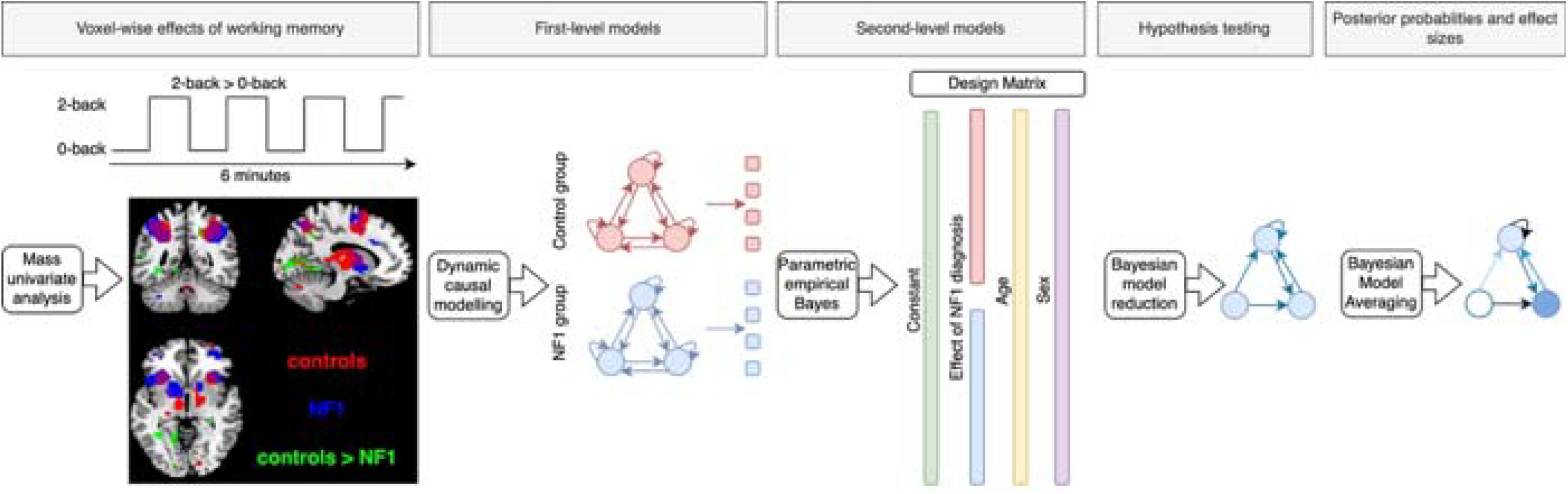
A visual summary of the fMRI analysis pipeline. In a summary, the mass univariate analysis was performed to identify effect of working memory in control and NF1 groups (2-back>0-back contrast), and then the areas where working memory evokes greater activity in controls than NF1 were identified (controls>NF1 contrast). Volumes showing these group effects were carried to first-level DCM. Group effects on effective connectivity were identified with parametric empirical Bayes approach. Connections that had no contribution to the model fit were removed with Bayesian model reduction. Finally, connection weights were averaged with Bayesian model averaging.

For each subject, a canonical hemodynamic response function was fitted for 0-back and 2-back conditions of N-back task and general linear model (GLM) was estimated. A high-pass filter with a cut-off at 128 seconds was applied to remove slow signal drifts. An autoregressive model (AR(1)) was fitted to estimate and remove serial correlations. Global masking threshold remained at default 0.8 value. The volume censoring regressors and 6 motion parameters were included as covariates in the model. One F-contrast of main effect of condition was defined, followed by a 2-back>0-back t-contrast.

We estimated the effect of working memory (2-back>0-back contrast) in each group. Then, from this contrast we created a union mask (uncorrected p-value 0.001). Within this union mask, we compared the effect of working memory across NF1 and control groups with whole brain mass univariate analysis (controls > NF1 group contrast). Age and sex were included in the design matrix as covariates of no interest.

### Volumes of Interest

Eight volumes of interest (VOIs) were defined in frontal and parietal lobes, based on existing meta-analysis of verbal and identity monitoring working memory (Emch et al., 2019; Owen et al., 2005). VOIs were defined as spheres with radius of 6mm, and they were additionally masked by the union mask of effect of working memory. This ensured that models reflect shared and unique patterns of neuronal activity related to working memory, rather than background activity. VOIs included bilateral superior parietal gyrus (SPG, MNI: -20, -56, 40; 20, -56, 50), bilateral ventrolateral prefrontal cortex (VLPFC, MNI: 32, 20, 4; -32, 24, 8), bilateral dorsolateral prefrontal cortex (DLPFC, MNI: -48, 16, 24; 48, 16, 24). Additionally, bilateral inferior parietal gyrus (IPG) was added based on peak working memory activation across all participants (MNI: -34, -44, 38; 36, -46, 40).

For every participant, the first eigenvariate of the timeseries of all voxels in each VOI was extracted and adjusted by F-contrast of main effects of the N-back task, which removed effects of motion and outlier scans from the timeseries.

### Dynamic Causal Models

Once the time series of VOIs were extracted, DCM for fMRI was used to estimate the effective connectivity for each participant (Friston et al., 2003). Each model was fully connected and composed of 40 intrinsic and modulatory connections between the 8 VOIs. Based on the prior literature, the driving input entered IPG (Ma et al., 2011).

Following the inversion of first level DCMs, we performed second level-analysis using the parametric empirical Bayes (PEB) approach (Friston et al., 2016;

Zeidman, Jafarian, Seghier, et al., 2019). This revealed the group mean connectivity shared across the control and NF1 groups, and the effect of NF1 diagnosis on connectivity. The PEB analysis was ran separately for A-matrix and B-matrix. We included age and sex as covariates of no interest and we mean-centred the design matrix.

Next, we investigated our hypothesis that the NF1 diagnosis would be associated with increased strength of inhibitory self-connections during working memory in NF1 group (intrinsic connectivity in B-matrix). Therefore, we sequentially “switched off” self-connections of every region and compared model evidence between the full and reduced (nested) models, which differed only in their priors. We used Bayesian Model Reduction (BMR) to estimate the free energy (an approximation of a model’s log-evidence) of reduced models and their corresponding parameters. This enabled Bayesian model comparison (BMC) by comparing the free energy of each model (Friston et al., 2016). Next, Bayesian model average (BMA) was used to obtain average of parameter estimates across models weighted by each model’s posterior probability (Penny et al., 2007). Posterior probability parameters of individual connections represent the effect sizes (Friston et al., 2016). The evidence level of the models is interpreted according to Kass and Raftery (1995). Finally, leave-one-out cross-validation (LOO-CV) was used to assess how well the estimated brain connectivity patterns could predict diagnosis status. At each fold, the PEB approach was repeated using all participants except one, to estimate group-level connectivity parameters for intrinsic connections identified via BMC. The diagnosis for the held-out participant was then predicted based on these estimated parameters. This process was repeated until each participant had been excluded once.

## Results

### Mass Univariate Analysis

The effects of working memory (2-back>0-back contrast) were evaluated separately for NF1 and control groups. Details of the local activation peaks are provided in Supplementary Tables 1 and 2 for controls and NF1, respectively. In both groups, working memory engagement was associated with bilateral activation in frontal regions, including superior, middle, and inferior frontal gyri. Additional activation was observed in the precentral gyri and supplementary motor area. In the parietal cortex, both groups showed activation in IPG and precuneus. Other regions showing significant working memory effect included left insula and left globus pallidus.

Supplementary Table 3 describes the local maxima of controls>NF1 contrast. Figure 3 illustrates the binarized mask of this contrast and the location of the VOIs. As hypothesized, working memory evoked greater activation in controls in the frontoparietal network, including bilateral SPG, bilateral VLPFC, and left DLPFC (all passing the uncorrected p-value threshold of 0.001). There was a relatively weak but significant difference in activation of left IPG, left supramarginal gyrus and right angular gyrus. Differences in activation also extended over bilateral insula, left middle cingulate cortex, left calcarine fissure, left middle temporal gyrus, and right cerebellum.

**Figure 3.**
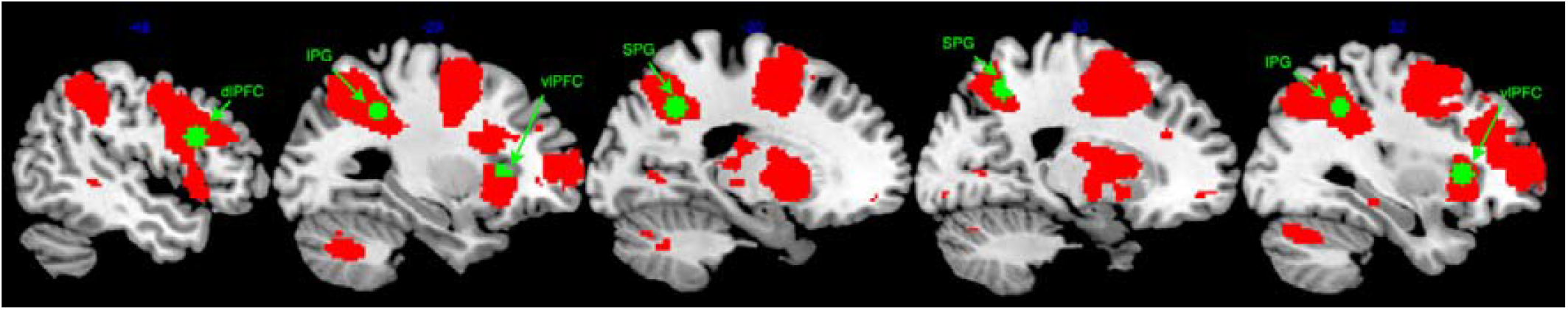
A binarized overlay of clusters generated by mass univariate analysis showing regions where the control group has greater working memory related activation than the NF1 group. Green voxels illustrate the VOIs. The sagittal slices have the following coordinates: -48, -29, -20, 20, 32.

### Effective connectivity Average connectivity

The PEB approach was used to estimate the shared and unique connections across groups based on the average connectivity of 0-back and 2-back conditions (A-matrix). Top graph in figure 4 illustrates common (i.e. shared) connection strength and bottom graph illustrates effect of NF1 diagnosis (whether it was associated with increase or decrease to connection strength). In this text, we focus on group differences that have strong evidence (posterior probability of including the connection in the model >0.95%). In NF1 group, relative to controls, there was stronger (i.e. more inhibitory) intrinsic connectivity in all regions of left hemisphere, and right vlPFC. Across hemispheres, parietal regions (both IPG and SPG) shared more excitatory cross-hemispheric connections to their contralateral counterparts. Focusing on left hemisphere, left dlPFC had less excitatory connection to SPG and more inhibitory connections to right dlPFC, but more excitatory connection to vlPFC. Left IPG projected less excitatory connection to SPG. In addition, left IPG had more inhibitory connection to vlPFC, but also IPG received more inhibitory connection from vlPFC. Finally, in right hemisphere, right dlPFC projected more inhibitory connections to IPG and SPG, but exchanged more excitatory connections with vlPFC. In parietal regions right SPG had a more excitatory connection to IPG.

**Figure 4.**
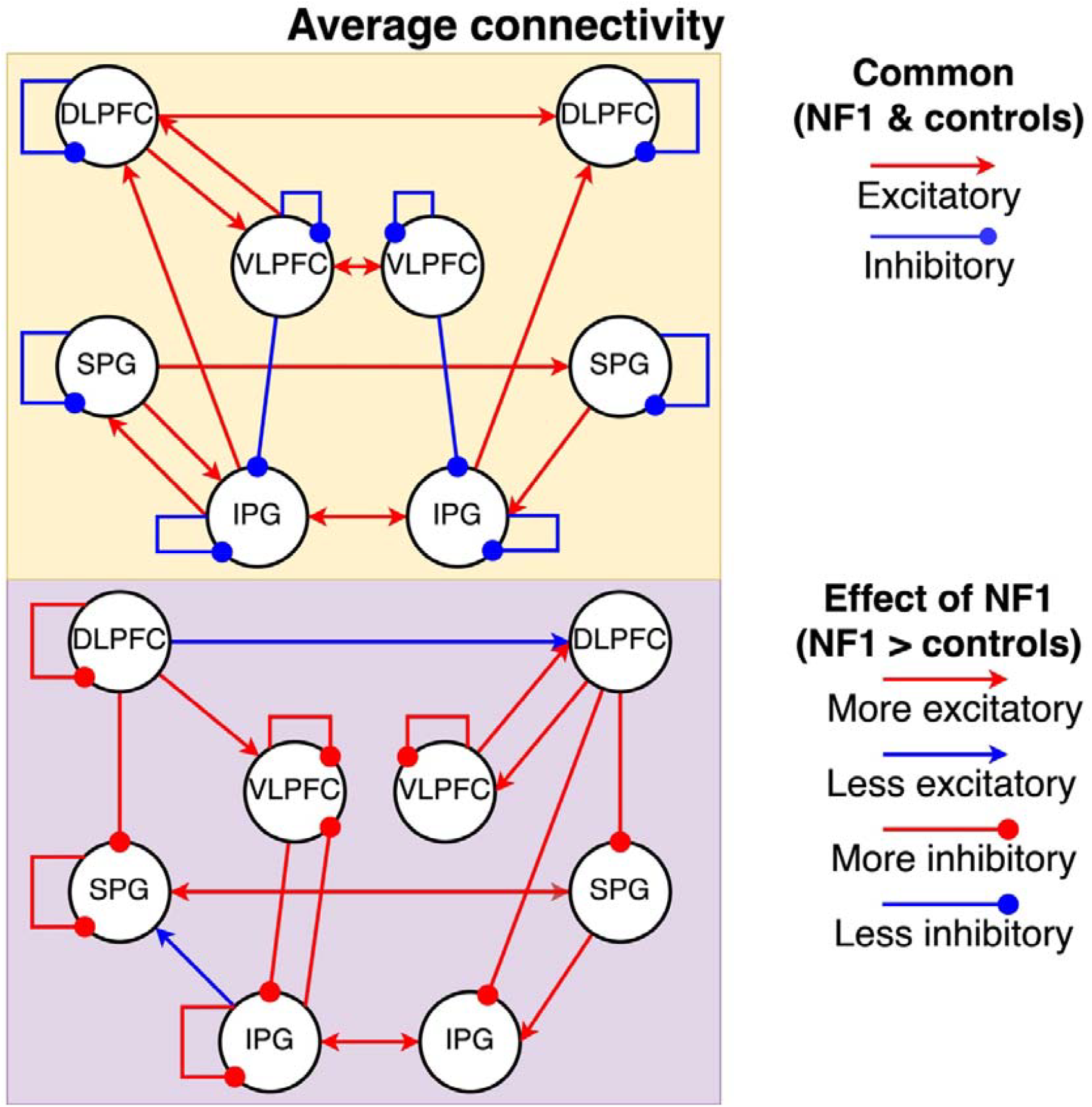
The average connectivity during 0-back and 2-back conditions of the N-back task. Top panel shows connections common to both groups; bottom panel shows group differences (NF1>controls). All displayed connections exceed a 95% posterior probability threshold. In the top panel red indicates excitatory and blue inhibitory influences. In the bottom panel, red connections are stronger in NF1 and blue weaker. For intrinsic (self) connections, “stronger” indicates greater self-inhibition.

### Modulatory connectivity

The PEB method was separately implemented to investigate the modulatory effect of working memory (B-matrix) (Figure 5). Again, we focus on strong evidence for group differences. In NF1 group, working memory evoked weaker (less inhibitory) intrinsic connectivity of left dlPFC. NF1 also had more inhibitory connections from left IPG to right IPG and left vlPFC, and from right dlPFC to right SPG. In NF1, there was a more excitatory connection from right to left SPG.

**Figure 5.**
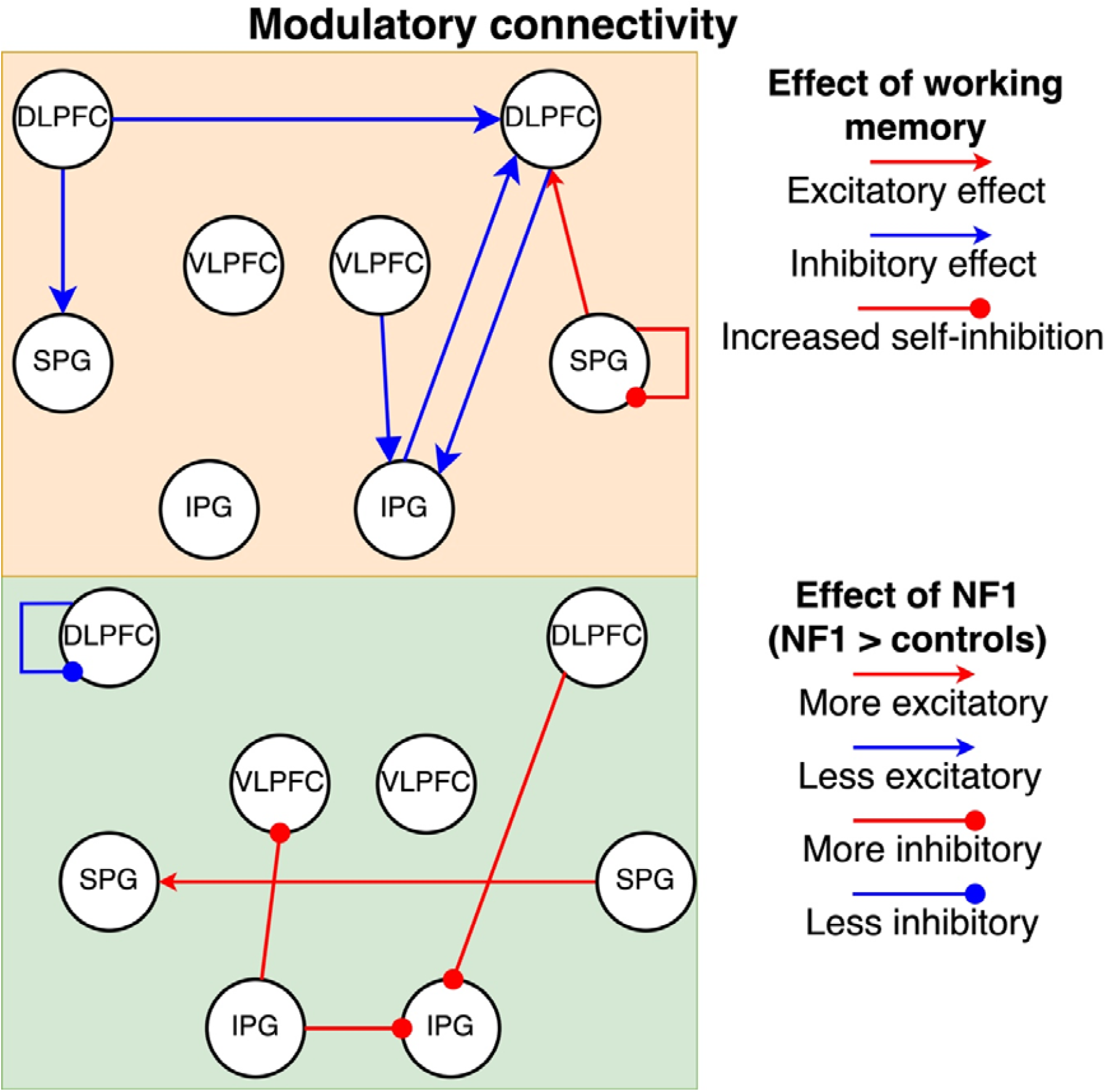
Modulatory effect of working memory on effective connectivity. Top panel shows connections common to both groups; bottom panel shows group differences (NF1>controls). All displayed connections exceed a 95% posterior probability threshold. In the top panels red indicates excitatory and blue inhibitory influences. In the bottom panel, red connections are stronger in NF1 and blue weaker. For intrinsic (self) connections, “stronger” indicates greater self-inhibition.

### Hypothesis testing

BMR was conducted to test the hypothesis that differences in extrinsic modulatory connections were driven by intrinsic connectivity. The winning model according to BMC had a posterior probability (*p*_post_) of 0.34%. It suggested that during working memory NF1 participants have stronger (i.e. more inhibitory) intrinsic connectivity of left vlPFC (*p*_post_ = 52%), and weaker intrinsic connectivity of right vlPFC (*p*_post_ = 52%) and left dlPFC (*p*_post_ = 89%). The parameters of these connections were weakly predictive of NF1 diagnosis; correlation coefficient between true and predictive scores was 0.19 (*p* = 0.055).

## Discussion

In this work, we investigated if NF1 participants have greater inhibitory activity in the frontoparietal network related to verbal working memory, compared to neurotypical controls. First, we observed that average intrinsic connectivity (i.e. average of 0-back and 2-back conditions) of left dlPFC, IPG, SPG and bilateral vlPFC was increased in NF1, suggesting reduced overall sensitivity of these regions to cognitive inputs. Second, in support of our hypothesis, using DCM analysis, we found that group differences during working memory are driven by more inhibitory intrinsic connectivity of left dlPFC and left vlPFC, and less inhibitory intrinsic connectivity of right vlPFC. During LOO-CV, these connectivity patterns showed a weak trend toward distinguishing between NF1 and control groups. Together, these findings suggests that NF1 participants have less sensitivity to task-related inputs and additionally may be less able to adapt to the task demands of working memory.

To study working memory differences, we selected frontal and parietal regions that showed activity related to working memory in both groups. A meta-analysis has related the activity of these regions to identity monitoring of verbal stimuli (Owen et al., 2005). In addition, these regions showed reduced BOLD activity in response to verbal working memory demands in NF1 participants, as compared to neurotypical controls. This finding of hypoactivity echoes similar findings from previous NF1 literature related to visuospatial working memory (Ibrahim et al., 2017; Shilyansky et al., 2010), and this suggests that hypoactivity may be a consistent feature across cognitive domains. We followed this pattern with DCM analysis, aiming to explore how the neuronal populations within these regions respond to cognitive inputs and specifically to working memory demands.

First, we focused on analysis of DCM’s A-matrix, which reflects the average connectivity during 0-back and 2-back conditions. This revealed that NF1 participants had stronger intrinsic average connectivity in left hemisphere. In DCM, intrinsic connectivity can be understood as the sensitivity of the region to inputs, where stronger intrinsic connectivity suggests more inhibition of input-related signals (Snyder et al., 2021). Therefore, our results suggest NF1 group has less sensitivity to the N-back task overall in the left hemisphere of frontoparietal network and right vlPFC. Due to this reduced sensitivity, NF1 group may have to recruit additional neural resources to perform the N-back task at the same level as neurotypical controls. This interpretation is consistent with neural efficiency hypothesis that neurodivergent individuals need to compensate/adapt to their condition through greater neural engagement for equivalent performance (Neubauer & Fink, 2009).

However, Poldrack (2015) raises an important point that observed differences may be indicative of different cognitive processes, different neural computations, differences in neural firing intensity or duration, or true metabolic efficiency differences. Here, our implementation of DCM becomes particularly useful in addressing this ambiguity because it also estimates modulatory connectivity, which here reflects the effect of working memory. Modulatory connectivity showed that working memory evoked less inhibitory intrinsic connectivity in left dlPFC and left vlPFC, but more inhibitory intrinsic connectivity in right vlPFC compared to controls. We speculate that this may occur due to high use of neural resources at baseline, which could limit the system’s capacity to adapt further to more demanding tasks.

Future research should investigate this ceiling effect by investigating if it can also be found in other executive functions, such as inhibitory control or task switching. Modulatory connectivity also revealed that during working memory, NF1 participants showed reduced inhibitory responses in left-lateralized prefrontal regions (left dlPFC and vlPFC), which suggests that NF1 participants have difficulty modulating their neural responses in response to working memory demands. This may indicate compromised phonological rehearsal process during working memory, because previous research indicates that there is left-lateralised specialisation of phonological rehearsal during verbal working memory (Crottaz-Herbette et al., 2004; Emch et al., 2019; Walter et al., 2003). The reduced inhibitory responses in left prefrontal regions may indicate reduced engagement of these phonological rehearsal processes in adolescents with NF1, which is consistent with previous reports of compromised phonological processing and memory in this population (Arnold et al., 2018; Baudou et al., 2020; Cutting & Levine, 2010).

We can further make sense of these differences in cognitive processing between NF1 and controls by considering the cognitive role of individual regions in the fronto-parietal network. We found that the right vlPFC had stronger inhibitory intrinsic connectivity in NF1 group than the control group. This region has been related to inhibitory control, attention regulation, and filtering of distractors (Aron et al., 2014; Hampshire et al., 2010; Shanmugan et al., 2016). More intrinsic inhibition of right vlPFC suggests that it is less sensitive to inputs, therefore it attenuates more stimuli. In context of this, we can hypothesize that the NF1 participants stayed more ‘on-line’ with the presented stimuli, and by extension it is possible that they committed fewer items to their memory. In contrast, left vlPFC and dlPFC were less inhibited (i.e. more driven by the inputs). Left vlPFC is known for its contribution to modulation of phonological processing and rehearsal (Owen et al., 2005; Walter et al., 2003). This suggests that instead of modulating the phonological inputs, the NF1 participants are more driven by the stimuli. Similarly, we found that left dlPFC was also less inhibited under working memory demands. Individual studies and meta-analysis show that that dlPFC is involved with executive function, working memory, cognitive control, and attention (Jimura et al., 2018; Kim et al., 2015; Osaka et al., 2003; Owen et al., 2005; Rottschy et al., 2012; Vartanian et al., 2013; Wager & Smith, 2003; Yaple et al., 2019). Specifically, Huey et al. (2016) proposed that left dlPFC is involved with monitoring of individual stimuli or events. Importantly, working memory deficits in individuals with NF1 have been linked to hypoactivity of left dlPFC (Ibrahim et al., 2017; Shilyansky et al., 2010). Here we further add to these results the finding that this hypoactivity is related to disrupted inhibitory activity in the region. Future research should focus on investigating these possibilities by correlating working memory performance to performance on tasks dedicated to measuring NF1 participants’ phonological processing, their ability to filter out distractor stimuli and their ability to commit novel items to memory.

The present work also carries important implications for research focusing on non-pharmacological interventions for NF1. Previous research demonstrated that non-invasive brain stimulation (NIBS) can be used to reduce GABA neurotransmitter concentration in left dlPFC (Garg et al., 2022). Our subsequent work employed the DCM method to understand how NIBS changed the dynamics of these brain regions (Litwińczuk et al., 2025). We showed that after NIBS, less dlPFC GABA was associated with less inhibition of left dlPFC throughout the N-back task (i.e. weaker intrinsic average connectivity). The present results from this investigation comparing DCM of NF1 group to control group add further insight to our understanding of the effects of NIBS; we now see that NIBS has addressed precisely the mechanisms underlying an atypical response in NF1 (i.e. increased intrinsic average connectivity). We also add the new and important finding that NF1 affects inhibitory activity in multiple regions of frontoparietal network (bilateral vlPFC and left dlPFC). These regions may constitute suitable stimulation sites in future NIBS research. In particular, it will be important to determine what NIBS protocols should be implemented (e.g. optimised electrode placements, use of transcranial direct current stimulation or transcranial alternating current stimulation).

It is important to understand this discussion in the context of methodological limitations of our analysis. The DCM analysis produces models of interactions between a set of brain regions. This means that other neuronal populations, outside the set of chosen VOIs also likely contribute to the dynamics of the regions we choose to study. In other words, the hypoactivity reported here is limited to the context of processes of frontoparietal networks. To illustrate, we explored in this analysis how working memory inputs enter, how they are maintained and manipulated, but we cannot comment on other processes such as inhibitory control, attentional processing, early visual processes or motor processing. Each of these components is a subject of its own study and offers opportunities for future research. Additionally, it is important to note that NF1 participants were not off their regular medications that may influence their cognitive processing. We did not include either medication status in the PEB analysis, nor comorbid diagnosis with over neurodevelopmental conditions.

Overall, this work has demonstrated that NF1 participants have increased average intrinsic connectivity during N-back task, suggesting increased inhibition and therefore reduced sensitivity to inputs. Additionally, NF1 participants showed reduced inhibitory responses in left-lateralized prefrontal regions during working memory demands but increased inhibitory activity in right vlPFC. These findings suggest that NF1 participants have less sensitivity to task-related inputs and additionally may be less able to adapt to the task demands of working memory.

## Acknowledgements

This research was supported by the NIHR Manchester Biomedical Research Centre (NIHR203308). ML was funded by the Office for Life Sciences and the National Institute for Health and Care Research (NIHR) Mental Health Translational Research Collaboration, hosted by the NIHR Oxford Health Biomedical Research Centre (NIHR203308). The views expressed are those of the author(s) and not necessarily those of the NIHR or the Department of Health and Social Care. The authors also wish to thank the study participants and their families. This work was also supported by the Neurofibromatosis Therapeutic Acceleration Program (NTAP) through a Francis Collins Scholarship to SG. NT is supported by Medical Research Council (MRC) (MR/X005267/1).

## References

Arnold, S. S., Payne, J. M., Lorenzo, J., North, K. N., & Barton, B. (2018). Preliteracy impairments in children with neurofibromatosis type 1. Developmental Medicine & Child Neurology, 60(7), 703–710. 10.1111/dmcn.13768

Aron, A. R., Robbins, T. W., & Poldrack, R. A. (2014). Inhibition and the right inferior frontal cortex: one decade on. Trends in Cognitive Sciences, 18(4), 177–185. 10.1016/j.tics.2013.12.003

Ashburner, J. (2007). A fast diffeomorphic image registration algorithm. NeuroImage, 38(1), 95–113. 10.1016/j.neuroimage.2007.07.007

Baddeley, A. (1992). Working memory.

Baddeley, A., Eysenck, M. W., & Anderson, M. C. (2014). Memory. Taylor & Francis Group. http://ebookcentral.proquest.com/lib/manchester/detail.action?docID=1843493

Bannai, H., Niwa, F., Sherwood, Mark W., Shrivastava, Amulya N., Arizono, M., Miyamoto, A., Sugiura, K., Lévi, S., Triller, A., & Mikoshiba, K. (2015). Bidirectional Control of Synaptic GABAAR Clustering by Glutamate and Calcium. Cell Reports, 13(12), 2768–2780. 10.1016/j.celrep.2015.12.002

Baudou, E., Nemmi, F., Biotteau, M., Maziero, S., Peran, P., & Chaix, Y. (2020). Can the Cognitive Phenotype in Neurofibromatosis Type 1 (NF1) Be Explained by Neuroimaging? A Review. Frontiers in Neurology, 10. 10.3389/fneur.2019.01373

Buratti, E., Karlsgodt, K. H., Rosser, T., Lutkenhoff, E. S., Cannon, T. D., Silva, A., & Bearden, C. E. (2012). Alterations in White Matter Microstructure in Neurofibromatosis-1. PLoS ONE, 7(10). 10.1371/journal.pone.0047854

Costa, R. M., Federov, N. B., Kogan, J. H., Murphy, G. G., Stern, J., Ohno, M., Kucherlapati, R., Jacks, T., & Silva, A. J. (2002). Mechanism for the learning deficits in a mouse model of neurofibromatosis type 1. Nature, 415(6871), 526–530. 10.1038/nature711

Crottaz-Herbette, S., Anagnoson, R. T., & Menon, V. (2004). Modality effects in verbal working memory: differential prefrontal and parietal responses to auditory and visual stimuli. NeuroImage, 21(1), 340–351. 10.1016/j.neuroimage.2003.09.019

Cutting, L. E., & Levine, T. M. (2010). Cognitive Profile of Children with Neurofibromatosis and Reading Disabilities. Child Neuropsychology, 16(5), 417–432. 10.1080/09297041003761985

Daston, M. M., & Ratner, N. (2005). Neurofibromin, a predominantly neuronal GTPase activating protein in the adult, is ubiquitously expressed during development. Developmental Dynamics, 195(3), 216–226. 10.1002/aja.1001950307

Emch, M., von Bastian, C. C., & Koch, K. (2019). Neural Correlates of Verbal Working Memory: An fMRI Meta-Analysis. Frontiers in Human Neuroscience, 13. 10.3389/fnhum.2019.00180

Evans, D. G., Howard, E., Giblin, C., Clancy, T., Spencer, H., Huson, S. M., & Lalloo, F. (2010). Birth incidence and prevalence of tumor-prone syndromes: Estimates from a UK family genetic register service. American Journal of Medical Genetics Part A, 152A(2), 327–332. 10.1002/ajmg.a.33139

Friston, K. J., Harrison, L., & Penny, W. (2003). Dynamic causal modelling. NeuroImage, 19(4), 1273–1302. 10.1016/s1053-8119(03)00202-7

Friston, K. J., Litvak, V., Oswal, A., Razi, A., Stephan, K. E., van Wijk, B. C. M., Ziegler, G., & Zeidman, P. (2016). Bayesian model reduction and empirical Bayes for group (DCM) studies. NeuroImage, 128, 413–431. 10.1016/j.neuroimage.2015.11.015

Garg, S., Green, J., Leadbitter, K., Emsley, R., Lehtonen, A., Evans, D. G., & Huson, S. M. (2013). Neurofibromatosis Type 1 and Autism Spectrum Disorder. Pediatrics, 132(6), e1642–e1648. 10.1542/peds.2013-1868

Garg, S., Lehtonen, A., Huson, S. M., Emsley, R., Trump, D., Evans, D. G., & Green, J. (2012). Autism and other psychiatric comorbidity in neurofibromatosis type 1: evidence from a population-based study. Developmental Medicine & Child Neurology, 55(2), 139–145. 10.1111/dmcn.12043

Garg, S., Williams, S., Jung, J., Pobric, G., Nandi, T., Lim, B., Vassallo, G., Green, J., Evans, D. G., Stagg, C. J., Parkes, L. M., & Stivaros, S. (2022). Non-invasive brain stimulation modulates GABAergic activity in neurofibromatosis 1. Scientific Reports, 12(1). 10.1038/s41598-022-21907-9

Gathercole, S. E., & Alloway, T. P. (2005). Practitioner Review: Short-term and working memory impairments in neurodevelopmental disorders: diagnosis and remedial support. Journal of Child Psychology and Psychiatry, 47(1), 4–15. 10.1111/j.1469-7610.2005.01446.x

Gutmann, D. H., Ferner, R. E., Listernick, R. H., Korf, B. R., Wolters, P. L., & Johnson, K. J. (2017). Neurofibromatosis type 1. Nature Reviews Disease Primers, 3(1). 10.1038/nrdp.2017.4

Halai, A. D., Welbourne, S. R., Embleton, K., & Parkes, L. M. (2014). A comparison of dual gradient-echo and spin-echo fMRI of the inferior temporal lobe. Human Brain Mapping, 35(8), 4118–4128. 10.1002/hbm.22463

Hampshire, A., Chamberlain, S. R., Monti, M. M., Duncan, J., & Owen, A. M. (2010). The role of the right inferior frontal gyrus: inhibition and attentional control. NeuroImage, 50(3), 1313–1319. 10.1016/j.neuroimage.2009.12.109

Huey, E. D., Krueger, F., & Grafman, J. (2016). Representations in the Human Prefrontal Cortex. Current Directions in Psychological Science, 15(4), 167–171. 10.1111/j.1467-8721.2006.00429.x

Ibrahim, A. F. A., Montojo, C. A., Haut, K. M., Karlsgodt, K. H., Hansen, L., Congdon, E., Rosser, T., Bilder, R. M., Silva, A. J., & Bearden, C. E. (2017). Spatial working memory in neurofibromatosis 1: Altered neural activity and functional connectivity. NeuroImage: Clinical, 15, 801–811. 10.1016/j.nicl.2017.06.032

Jimura, K., Chushak, M. S., Westbrook, A., & Braver, T. S. (2018). Intertemporal Decision-Making Involves Prefrontal Control Mechanisms Associated with Working Memory. Cerebral Cortex, 28(4), 1105–1116. 10.1093/cercor/bhx015

Kass, R. E., & Raftery, A. E. (1995). Bayes Factors. Journal of the American Statistical Association, 90(430), 773–795. 10.1080/01621459.1995.10476572

Kim, C., Kroger, J. K., Calhoun, V. D., & Clark, V. P. (2015). The role of the frontopolar cortex in manipulation of integrated information in working memory. Neuroscience Letters, 595, 25–29. 10.1016/j.neulet.2015.03.044

Lehtonen, A., Howie, E., Trump, D., & Huson, S. M. (2012). Behaviour in children with neurofibromatosis type 1: cognition, executive function, attention, emotion, and social competence. Developmental Medicine & Child Neurology, 55(2), 111–125. 10.1111/j.1469-8749.2012.04399.x

Litwińczuk, M. C., Garg, S., Williams, S. R., Green, J., Lea-Carnall, C., & Trujillo-Barreto, N. J. (2025). Non-invasive brain stimulation reorganises effective connectivity during a working memory task in individuals with Neurofibromatosis Type 1. NeuroImage: Reports, 5(2). 10.1016/j.ynirp.2025.100258

Ma, L., Steinberg, J. L., Hasan, K. M., Narayana, P. A., Kramer, L. A., & Moeller, F. G. (2011). Working memory load modulation of parieto-frontal connections: Evidence from dynamic causal modeling. Human Brain Mapping, 33(8), 1850–1867. 10.1002/hbm.21329

Maffei, A., Charrier, C., Caiati, M. D., Barberis, A., Mahadevan, V., Woodin, M. A., & Tyagarajan, S. K. (2017). Emerging Mechanisms Underlying Dynamics of GABAergic Synapses. The Journal of Neuroscience, 37(45), 10792–10799. 10.1523/jneurosci.1824-17.2017

Matson, J. L., Rieske, R. D., & Williams, L. W. (2013). The relationship between autism spectrum disorders and attention-deficit/hyperactivity disorder: An overview. Research in Developmental Disabilities, 34(9), 2475–2484. 10.1016/j.ridd.2013.05.021

Metkus, J. D., Blanco, T., Mohan, A., Oh, A., Robinson, C., & Bhattacharya, S. (2024). Excitation–inhibition balance in diseases of the brain: Role of NMDA and GABA receptors. In A Review on Diverse Neurological Disorders (pp. 353–383). 10.1016/b978-0-323-95735-9.00021-8

Neubauer, A. C., & Fink, A. (2009). Intelligence and neural efficiency: Measures of brain activation versus measures of functional connectivity in the brain. Intelligence, 37(2), 223–229. 10.1016/j.intell.2008.10.008

North, K., Gutmann, D. H., & International Child Neurology, A. (1997). Neurofibromatosis type 1 in childhood.

Osaka, M., Osaka, N., Kondo, H., Morishita, M., Fukuyama, H., Aso, T., & Shibasaki, H. (2003). The neural basis of individual differences in working memory capacity: an fMRI study. NeuroImage, 18(3), 789–797. 10.1016/s1053-8119(02)00032-0

Owen, A. M., McMillan, K. M., Laird, A. R., & Bullmore, E. (2005). N-back working memory paradigm: A meta-analysis of normative functional neuroimaging studies. Human Brain Mapping, 25(1), 46–59. 10.1002/hbm.20131

Payne, J. M., Moharir, M. D., Webster, R., & North, K. N. (2010). Brain structure and function in neurofibromatosis type 1: current concepts and future directions. Journal of Neurology, Neurosurgery & Psychiatry, 81(3), 304–309. 10.1136/jnnp.2009.179630

Penny, W. D., Mattout, J., & Trujillo-Barreto, N. (2007). Bayesian model selection and averaging. In Statistical Parametric Mapping (pp. 454–467). 10.1016/b978-012372560-8/50035-8

Petroff, O. A. C. (2016). Book Review: GABA and Glutamate in the Human Brain. The Neuroscientist, 8(6), 562–573. 10.1177/1073858402238515

Pobric, G., Taylor, J. R., Ramalingam, H. M., Pye, E., Robinson, L., Vassallo, G., Jung, J., Bhandary, M., Szumanska-Ryt, K., Theodosiou, L., Evans, D. G., Eelloo, J., Burkitt-Wright, E., Hulleman, J., Green, J., & Garg, S. (2021). Cognitive and Electrophysiological Correlates of Working Memory Impairments in Neurofibromatosis Type 1. Journal of Autism and Developmental Disorders, 52(4), 1478–1494. 10.1007/s10803-021-05043-3

Poldrack, R. A. (2015). Is “efficiency” a useful concept in cognitive neuroscience? Developmental Cognitive Neuroscience, 11, 12–17. 10.1016/j.dcn.2014.06.001

Rottschy, C., Langner, R., Dogan, I., Reetz, K., Laird, A. R., Schulz, J. B., Fox, P. T., & Eickhoff, S. B. (2012). Modelling neural correlates of working memory: A coordinate-based meta-analysis. NeuroImage, 60(1), 830–846. 10.1016/j.neuroimage.2011.11.050

Sawyer, C., Green, J., Lim, B., Pobric, G., Jung, J., Vassallo, G., Evans, D. G., Stagg, C. J., Parkes, L. M., Stivaros, S., Muhlert, N., & Garg, S. (2022). Neuroanatomical correlates of working memory performance in Neurofibromatosis 1. Cerebral Cortex Communications, 3(2). 10.1093/texcom/tgac021

Sears, S. M. S., & Hewett, S. J. (2021). Influence of glutamate and GABA transport on brain excitatory/inhibitory balance. Experimental Biology and Medicine, 246(9), 1069–1083. 10.1177/1535370221989263

Shanmugan, S., Wolf, D. H., Calkins, M. E., Moore, T. M., Ruparel, K., Hopson, R. D., Vandekar, S. N., Roalf, D. R., Elliott, M. A., Jackson, C., Gennatas, E. D., Leibenluft, E., Pine, D. S., Shinohara, R. T., Hakonarson, H., Gur, R. C., Gur, R. E., & Satterthwaite, T. D. (2016). Common and Dissociable Mechanisms of Executive System Dysfunction Across Psychiatric Disorders in Youth. American Journal of Psychiatry, 173(5), 517–526. 10.1176/appi.ajp.2015.15060725

Shilyansky, C., Karlsgodt, K. H., Cummings, D. M., Sidiropoulou, K., Hardt, M., James, A. S., Ehninger, D., Bearden, C. E., Poirazi, P., Jentsch, J. D., Cannon, T. D., Levine, M. S., & Silva, A. J. (2010). Neurofibromin regulates corticostriatal inhibitory networks during working memory performance. Proceedings of the National Academy of Sciences, 107(29), 13141–13146. 10.1073/pnas.1004829107

Snyder, A. D., Ma, L., Steinberg, J. L., Woisard, K., & Moeller, F. G. (2021). Dynamic Causal Modeling Self-Connectivity Findings in the Functional Magnetic Resonance Imaging Neuropsychiatric Literature [Mini Review]. Frontiers in Neuroscience, 15. 10.3389/fnins.2021.636273

Sohal, V. S., & Rubenstein, J. L. R. (2019). Excitation-inhibition balance as a framework for investigating mechanisms in neuropsychiatric disorders. Molecular Psychiatry, 24(9), 1248–1257. 10.1038/s41380-019-0426-0

van der Meer, J. M. J., Oerlemans, A. M., van Steijn, D. J., Lappenschaar, M. G. A., de Sonneville, L. M. J., Buitelaar, J. K., & Rommelse, N. N. J. (2012). Are Autism Spectrum Disorder and Attention-Deficit/Hyperactivity Disorder Different Manifestations of One Overarching Disorder? Cognitive and Symptom Evidence From a Clinical and Population-Based Sample. Journal of the American Academy of Child & Adolescent Psychiatry, 51(11), 1160–1172.e1163. 10.1016/j.jaac.2012.08.024

van Lier, M., Saiepour, M. H., Kole, K., Cheyne, J. E., Zabouri, N., Blok, T., Qin, Y., Ruimschotel, E., Heimel, J. A., Lohmann, C., & Levelt, C. N. (2020). Disruption of Critical Period Plasticity in a Mouse Model of Neurofibromatosis Type 1. The Journal of Neuroscience, 40(28), 5495–5509. 10.1523/jneurosci.2235-19.2020

Vartanian, O., Jobidon, M. E., Bouak, F., Nakashima, A., Smith, I., Lam, Q., & Cheung, B. (2013). Working memory training is associated with lower prefrontal cortex activation in a divergent thinking task. Neuroscience, 236, 186–194. 10.1016/j.neuroscience.2012.12.060

Wager, T. D., & Smith, E. E. (2003). Neuroimaging studies of working memory. Cognitive, Affective, & Behavioral Neuroscience, 3(4), 255–274. 10.3758/cabn.3.4.255

Walter, H., Bretschneider, V., Gron, G., Zurowski, B., Wunderlich, A., Tomczak, R., & Spitzer, M. (2003). Evidence for Quantitative Domain Dominance for Verbal and Spatial Working Memory in Frontal and Parietal Cortex. Cortex, 39(4-5), 897–911. 10.1016/s0010-9452(08)70869-4

Yaple, Z. A., Stevens, W. D., & Arsalidou, M. (2019). Meta-analyses of the n-back working memory task: fMRI evidence of age-related changes in prefrontal cortex involvement across the adult lifespan. NeuroImage, 196, 16–31. 10.1016/j.neuroimage.2019.03.074

Zeidman, P., Jafarian, A., Corbin, N., Seghier, M. L., Razi, A., Price, C. J., & Friston, K. J. (2019). A guide to group effective connectivity analysis, part 1: First level analysis with DCM for fMRI. NeuroImage, 200, 174–190. 10.1016/j.neuroimage.2019.06.031

Zeidman, P., Jafarian, A., Seghier, M. L., Litvak, V., Cagnan, H., Price, C. J., & Friston, K. J. (2019). A guide to group effective connectivity analysis, part 2: Second level analysis with PEB. NeuroImage, 200, 12–25. 10.1016/j.neuroimage.2019.06.032

