## Supplemental Material 1 for "Disrupted Frontoparietal Dynamics in Neurofibromatosis Type 1: Reduced Sensitivity and Atypical Modulation During Working Memory"

**Supplementary Material 1**

| Supplementary Table 1 Whole brain mass univariate analysis: brain regions showing significant effects of working memory in neurotypical controls. Statistical threshold was set at family wise error corrected *p* < 0.05. Region labels were generated with Automated Anatomical Labelling 3 (Rolls et al., 2020). | | | | | | | | | | |
| --- | --- | --- | --- | --- | --- | --- | --- | --- | --- | --- |
| cluster | | | peak | | | | MNI | | | Label |
| p (FWE) | p (FDR) | cluster size (k) | p (FWE) | p (FDR) | T-value | Z-value | x | y | z |  |
| 0 | 0 | 272 | 0 | 0.065 | 10.12 | 6.19 | 42 | -34 | 38 | Supramarginal gyrus |
|  |  |  | 0.002 | 0.21 | 8.25 | 5.57 | 28 | -56 | 40 | Angular gyrus |
|  |  |  | 0.005 | 0.231 | 7.7 | 5.36 | 32 | -52 | 46 | Inferior parietal gyrus |
| 0 | 0 | 256 | 0 | 0.15 | 8.98 | 5.83 | -22 | 0 | 44 | Superior frontal gyrus |
|  |  |  | 0.003 | 0.215 | 7.97 | 5.47 | -20 | 4 | 66 | Superior frontal gyrus, |
|  |  |  | 0.003 | 0.215 | 7.89 | 5.43 | -26 | 2 | 56 | Middle frontal gyrus |
| 0 | 0 | 57 | 0.001 | 0.15 | 8.86 | 5.79 | -2 | 6 | 58 | Supplementary motor area |
| 0 | 0 | 312 | 0.001 | 0.15 | 8.77 | 5.76 | -28 | -52 | 38 | Inferior parietal gyrus |
|  |  |  | 0.002 | 0.215 | 8.04 | 5.49 | -34 | -52 | 50 | Inferior parietal gyrus |
|  |  |  | 0.004 | 0.231 | 7.71 | 5.37 | -30 | -44 | 38 | Inferior parietal gyrus |
| 0 | 0 | 295 | 0.001 | 0.183 | 8.52 | 5.67 | 26 | 2 | 50 | Superior parietal gyrus |
|  |  |  | 0.002 | 0.21 | 8.26 | 5.58 | 26 | 0 | 62 | Superior parietal gyrus |
|  |  |  | 0.011 | 0.335 | 7.2 | 5.16 | 22 | 12 | 56 | Superior parietal gyrus |
| 0 | 0 | 74 | 0.002 | 0.215 | 8.06 | 5.5 | -32 | 18 | -2 | Insula |
| 0 | 0.008 | 30 | 0.003 | 0.215 | 7.9 | 5.44 | 18 | -62 | 56 | Superior parietal gyrus |
| 0 | 0 | 93 | 0.003 | 0.215 | 7.87 | 5.43 | 6 | 14 | 48 | Supplementary motor area |
|  |  |  | 0.006 | 0.267 | 7.57 | 5.31 | -2 | 14 | 46 | Supplementary motor area |
| 0 | 0.02 | 21 | 0.008 | 0.296 | 7.36 | 5.23 | 32 | 18 | -2 | Lenticular nucleus, Putamen |
| 0.004 | 0.141 | 8 | 0.009 | 0.296 | 7.35 | 5.22 | 46 | -38 | 54 | Inferior parietal gyrus |
| 0 | 0.018 | 23 | 0.009 | 0.314 | 7.3 | 5.2 | 18 | 0 | 12 | Caudate nucleus |
| 0 | 0.019 | 22 | 0.011 | 0.335 | 7.23 | 5.17 | -42 | -4 | 28 | Precentral gyrus |
| 0.001 | 0.042 | 15 | 0.011 | 0.335 | 7.2 | 5.16 | 46 | 26 | 28 | Inferior frontal gyrus, triangular part |
| 0.001 | 0.033 | 17 | 0.018 | 0.471 | 6.95 | 5.05 | 8 | -64 | 48 | Precuneus |
| 0.019 | 0.414 | 2 | 0.019 | 0.471 | 6.93 | 5.04 | 18 | 10 | 12 | Caudate nucleus |
| 0.008 | 0.255 | 5 | 0.024 | 0.589 | 6.78 | 4.98 | -14 | 0 | 6 | Lenticular nucleus, Pallidum |
| 0.019 | 0.414 | 2 | 0.031 | 0.717 | 6.65 | 4.92 | 34 | -68 | 40 | Occipital |
| 0.014 | 0.401 | 3 | 0.032 | 0.723 | 6.63 | 4.91 | 22 | 10 | 42 | Superior frontal gyrus |
| 0.019 | 0.414 | 2 | 0.034 | 0.732 | 6.61 | 4.9 | -36 | 58 | 8 | Middle frontal gyrus |
| 0.019 | 0.414 | 2 | 0.035 | 0.749 | 6.59 | 4.89 | -48 | 30 | 28 | Inferior frontal gyrus, triangular part |
| 0.019 | 0.414 | 2 | 0.039 | 0.818 | 6.53 | 4.86 | 36 | -70 | -28 | Crus I of cerebellar hemisphere |
| 0.027 | 0.541 | 1 | 0.045 | 0.917 | 6.46 | 4.83 | -34 | 18 | 22 | Inferior frontal gyrus, triangular part |
| 0.027 | 0.541 | 1 | 0.049 | 0.984 | 6.41 | 4.81 | -14 | -6 | 16 | Caudate nucleus |

| Supplementary Table 2 Whole brain mass univariate analysis: brain regions showing significant effects of working memory in NF1 patients. Statistical threshold was set at family wise error corrected *p* < 0.05. Region labels were generated with Automated Anatomical Labelling 3 (Rolls et al., 2020). | | | | | | | | | | |
| --- | --- | --- | --- | --- | --- | --- | --- | --- | --- | --- |
| cluster | | | peak | | | | MNI | | | Label |
| p (FWE) | p (FDR) | cluster size (k) | p (FWE) | p (FDR) | T-value | Z-value | x | y | z |  |
| 0 | 0 | 505 | 0 | 0 | 10.18 | 7.11 | 28 | 0 | 50 | Middle frontal gyrus |
|  |  |  | 0 | 0.019 | 7.65 | 5.97 | 32 | 2 | 60 | Superior frontal gyrus, dorsolateral |
| 0 | 0 | 858 | 0 | 0 | 9.9 | 7 | -2 | 12 | 50 | Supplementary motor area |
|  |  |  | 0 | 0.029 | 7.31 | 5.79 | 6 | 26 | 30 | Middle cingulate & paracingulate gyri |
|  |  |  | 0.001 | 0.032 | 7.1 | 5.68 | 0 | 26 | 40 | Superior frontal gyrus, medial |
| 0 | 0 | 615 | 0 | 0.011 | 8.15 | 6.22 | -36 | -44 | 38 | Inferior parietal gyrus |
|  |  |  | 0 | 0.016 | 7.92 | 6.1 | -42 | -50 | 46 | Inferior parietal gyrus |
|  |  |  | 0 | 0.022 | 7.47 | 5.88 | -30 | -58 | 48 | Inferior parietal gyrus |
| 0 | 0 | 832 | 0 | 0.019 | 7.72 | 6.01 | 42 | -44 | 50 | Inferior parietal gyrus |
|  |  |  | 0 | 0.029 | 7.3 | 5.79 | 12 | -70 | 54 | Precuneus |
|  |  |  | 0.001 | 0.03 | 7.21 | 5.74 | 38 | -46 | 42 | Inferior parietal gyrus |
| 0 | 0 | 113 | 0 | 0.019 | 7.67 | 5.98 | -26 | -60 | -36 | Lobule VI of cerebellar hemisphere |
|  |  |  | 0.003 | 0.104 | 6.55 | 5.37 | -40 | -70 | -34 | Crus I of cerebellar hemisphere |
| 0 | 0 | 184 | 0 | 0.022 | 7.47 | 5.88 | -46 | 18 | 36 | Middle frontal gyrus |
|  |  |  | 0.001 | 0.03 | 7.21 | 5.74 | -42 | 24 | 30 | Inferior frontal gyrus, triangular part |
|  |  |  | 0.001 | 0.031 | 7.13 | 5.69 | -46 | 30 | 34 | Middle frontal gyrus |
| 0 | 0 | 242 | 0.001 | 0.03 | 7.21 | 5.74 | -24 | 0 | 54 | Superior frontal gyrus, dorsolateral |
|  |  |  | 0.004 | 0.11 | 6.52 | 5.35 | -28 | 6 | 60 | Middle frontal gyrus |
| 0 | 0 | 131 | 0.001 | 0.031 | 7.14 | 5.7 | -46 | 16 | 0 | Inferior frontal gyrus, opercular part |
|  |  |  | 0.001 | 0.044 | 6.95 | 5.6 | -28 | 22 | -4 | Insula |
|  |  |  | 0.014 | 0.326 | 6.03 | 5.05 | -42 | 20 | -6 | Inferior frontal gyrus, orbital part |
| 0 | 0.002 | 47 | 0.001 | 0.031 | 7.14 | 5.7 | -34 | -2 | 38 | Precentral gyrus |
| 0 | 0 | 131 | 0.001 | 0.044 | 6.97 | 5.6 | 40 | 28 | 32 | Middle frontal gyrus |
| 0 | 0 | 67 | 0.002 | 0.066 | 6.75 | 5.48 | 34 | 20 | -6 | Insula |
| 0.003 | 0.071 | 14 | 0.004 | 0.123 | 6.45 | 5.31 | 50 | 8 | 36 | Precentral gyrus |
| 0.004 | 0.103 | 11 | 0.013 | 0.317 | 6.05 | 5.07 | -20 | 2 | 0 | Lenticular nucleus, Pallidum |
| 0.011 | 0.27 | 5 | 0.029 | 0.619 | 5.75 | 4.88 | 36 | 52 | 2 | Middle frontal gyrus |
| 0.017 | 0.388 | 3 | 0.03 | 0.62 | 5.74 | 4.87 | -10 | 2 | -2 | Lenticular nucleus, Pallidum |
| 0.022 | 0.469 | 2 | 0.034 | 0.69 | 5.69 | 4.84 | 26 | 46 | -14 | Anterior orbital gyrus |
| 0.03 | 0.598 | 1 | 0.044 | 0.878 | 5.59 | 4.78 | -18 | 6 | 10 | Lenticular nucleus, Putamen |

| Supplementary Table 3 Whole brain mass univariate analysis: brain regions showing greater working memory related activation in neurotypical controls than NF1 patients. Peak statistical threshold was uncorrected (*p* < 0.001). Region labels were generated with Automated Anatomical Labelling 3 (Rolls et al., 2020). | | | | | | | | | | |
| --- | --- | --- | --- | --- | --- | --- | --- | --- | --- | --- |
| cluster | | | peak | | | | MNI | | | Label |
| p (FWE) | p (FDR) | cluster size (k) | p (FWE) | p (FDR) | T-value | Z-value | x | y | z |  |
| 0.024 | 0.029 | 212 | 0.011 | 0.06 | 5.6 | 5.04 | -18 | -58 | 40 | Middle occipital gyrus |
| 0.009 | 0.016 | 265 | 0.082 | 0.233 | 4.97 | 4.56 | 20 | -56 | 50 | Superior parietal gyrus |
|  |  |  | 0.519 | 0.638 | 4.26 | 3.98 | 10 | -64 | 48 | Precuneus |
| 0.088 | 0.08 | 146 | 0.161 | 0.319 | 4.74 | 4.38 | -36 | 26 | 10 | Inferior frontal gyrus, triangular part |
|  |  |  | 0.449 | 0.638 | 4.33 | 4.04 | -34 | 18 | 12 | Insula |
|  |  |  | 0.609 | 0.638 | 4.17 | 3.91 | -26 | 24 | 0 | Insula |
| 0.001 | 0.002 | 435 | 0.573 | 0.638 | 4.2 | 3.94 | -16 | -68 | 6 | Calcarine fissure and surrounding cortex |
|  |  |  | 0.819 | 0.734 | 3.95 | 3.72 | -14 | -52 | -4 | Lingual gyrus |
|  |  |  | 0.821 | 0.734 | 3.94 | 3.72 | -14 | -82 | 2 | Lingual gyrus |
| 0.811 | 0.646 | 34 | 0.584 | 0.638 | 4.19 | 3.93 | -10 | 10 | 38 | Middle cingulate & paracingulate gyri |
| 0.533 | 0.402 | 59 | 0.71 | 0.734 | 4.07 | 3.82 | 22 | -6 | 40 | Middle cingulate & paracingulate gyri |
| 0.901 | 0.698 | 25 | 0.755 | 0.734 | 4.02 | 3.79 | -54 | -48 | 14 | Middle temporal gyrus |
| 0.974 | 0.704 | 14 | 0.905 | 0.734 | 3.82 | 3.62 | -24 | 46 | 12 | Middle frontal gyrus |
| 0.553 | 0.402 | 57 | 0.935 | 0.734 | 3.77 | 3.57 | -38 | 4 | 22 | Inferior frontal gyrus, opercular part |
| 0.883 | 0.698 | 27 | 0.947 | 0.734 | 3.74 | 3.54 | 26 | -58 | 2 | Lingual gyrus |
| 0.947 | 0.704 | 19 | 0.961 | 0.734 | 3.7 | 3.51 | 30 | -32 | -6 | Hippocampus |
| 0.991 | 0.746 | 9 | 0.963 | 0.734 | 3.69 | 3.5 | 32 | -42 | 24 | Angular gyrus |
| 0.91 | 0.698 | 24 | 0.965 | 0.734 | 3.68 | 3.5 | 62 | -38 | 18 | Superior temporal gyrus |
| 0.249 | 0.2 | 96 | 0.973 | 0.734 | 3.65 | 3.47 | 22 | -74 | -28 | Crus I of cerebellar hemisphere |
|  |  |  | 0.983 | 0.734 | 3.6 | 3.43 | 30 | -70 | -26 | Lobule VI of cerebellar hemisphere |
| 0.986 | 0.704 | 11 | 0.974 | 0.734 | 3.65 | 3.47 | 32 | 26 | 8 | Insula |
| 0.974 | 0.704 | 14 | 0.98 | 0.734 | 3.62 | 3.44 | -48 | 16 | 24 | Inferior frontal gyrus, triangular part |
| 0.979 | 0.704 | 13 | 0.981 | 0.734 | 3.62 | 3.44 | 16 | -72 | 8 | Calcarine fissure and surrounding cortex |
| 0.959 | 0.704 | 17 | 0.986 | 0.734 | 3.58 | 3.41 | -48 | -36 | 30 | Supramarginal gyrus |
| 0.979 | 0.704 | 13 | 0.988 | 0.734 | 3.57 | 3.4 | -26 | 6 | 44 | Middle frontal gyrus |
| 0.778 | 0.646 | 37 | 0.989 | 0.734 | 3.56 | 3.39 | 16 | -94 | 0 | Calcarine fissure and surrounding cortex |
|  |  |  | 0.994 | 0.754 | 3.51 | 3.35 | 16 | -86 | 4 | Calcarine fissure and surrounding cortex |
| 0.954 | 0.704 | 18 | 0.99 | 0.734 | 3.55 | 3.38 | 20 | -90 | -8 | Lingual gyrus |
| 0.996 | 0.813 | 6 | 0.995 | 0.762 | 3.49 | 3.33 | 18 | 12 | 58 | Superior frontal gyrus, dorsolateral |
| 0.986 | 0.704 | 11 | 0.997 | 0.804 | 3.45 | 3.29 | 48 | -38 | 24 | Supramarginal gyrus |
| 1 | 0.845 | 1 | 0.999 | 0.931 | 3.35 | 3.21 | -12 | 10 | 56 | Superior frontal gyrus, dorsolateral |
| 0.995 | 0.798 | 7 | 0.999 | 0.931 | 3.35 | 3.2 | -28 | 2 | 20 | Supramarginal gyrus |
| 1 | 0.845 | 1 | 0.999 | 0.938 | 3.34 | 3.19 | -54 | -6 | 44 | Superior frontal gyrus, dorsolateral |
| 0.999 | 0.845 | 3 | 1 | 0.963 | 3.31 | 3.17 | 4 | -54 | -32 | Insula |
| 0.999 | 0.845 | 2 | 1 | 0.974 | 3.29 | 3.15 | -36 | 6 | 56 | Postcentral gyrus |
| 1 | 0.845 | 1 | 1 | 0.974 | 3.28 | 3.14 | 34 | -48 | 50 | Lobule IX of vermis |
| 1 | 0.845 | 1 | 1 | 0.974 | 3.27 | 3.13 | 4 | -66 | -20 | Middle frontal gyrus |
| 1 | 0.845 | 1 | 1 | 0.974 | 3.25 | 3.12 | -32 | -38 | 32 | Inferior parietal gyrus, excluding supramarginal and angular gyri |
| 0.999 | 0.845 | 3 | 1 | 0.974 | 3.24 | 3.1 | -8 | -66 | 44 | Precuneus |
| 1 | 0.845 | 1 | 1 | 0.974 | 3.24 | 3.1 | 22 | -32 | 8 | Hippocampus |
| 0.999 | 0.845 | 2 | 1 | 0.974 | 3.23 | 3.1 | -34 | -78 | 18 | Middle occipital gyrus |

Reference

Rolls, E. T., Huang, C.-C., Lin, C.-P., Feng, J., & Joliot, M. (2020). Automated anatomical labelling atlas 3. *NeuroImage*, *206*. <https://doi.org/10.1016/j.neuroimage.2019.116189>
